# Automated Separation of Diffusely Abnormal White Matter from Focal White Matter Lesions on MRI in Multiple Sclerosis

**DOI:** 10.1101/727818

**Authors:** Josefina Maranzano, Mahsa Dadar, Douglas L. Arnold, D. Louis Collins, Sridar Narayanan

## Abstract

**Background:** Previous histopathology and MRI studies have addressed the differences between focal white matter lesions (FWML) and diffusely abnormal white matter (DAWM) in multiple sclerosis (MS). These two categories of white matter T2-weighted (T2w) hyperintensity show different degrees of demyelination, axonal loss and immune cell density on histopathology, potentially offering distinct correlations with symptoms.

**Objectives:** 1) To automate the separation of FWML and DAWM using T2w MRI intensity thresholds and to investigate their differences in magnetization transfer ratios (MTR), which are sensitive to myelin content; 2) to correlate MTR values in FWML and DAWM with normalized signal intensity values on fluid attenuated inversion recovery (FLAIR), T2w, and T1-weighted (T1w) contrasts, as well as with the ratio of T2w/T1w normalized values, in order to determine whether these normalized intensities can be used as myelin-sensitive markers when MTR is not available.

**Methods:** Using similar 3T MRI protocols, 2 MS cohorts of 20 participants each were scanned in 2 centers, including: 3D T1w (MP2RAGE), 3D FLAIR, 2D T2w, and 3D magnetization-transfer (MT) contrasts. We used the first dataset to develop an automated technique to separate FWML from DAWM and the second one to validate the automation of the technique. We applied the automatic thresholds to both datasets and assessed the overlap of the manual and the automated masks using Dice kappa. We also assessed differences in mean MTR values between NAWM, DAWM and FWML, using manually and automatically derived masks in both datasets. Finally, we used the mean intensity of manually-traced areas of NAWM on T2w images as the normalization factor for each MRI contrast, and compared these with the normalized-intensity values obtained using automated NAWM (A-NAWM) masks as the normalization factor. Paired t-tests assessed the MTR differences across tissue types. Wilcoxon Signed Rank test and paired t-tests assessed FWML and DAWM differences in manual and automated derived volumes. Pearson correlations assessed the relationship between MTR and normalized intensity values in the manual and automated derived masks.

**Results:** The mean Dice-kappa values for dataset 1 were: 0.8 for DAWM masks and 0.7 for FWML masks. In dataset 2, mean dice-kappa values were: 0.8 for DAWM and 0.8 for FWML. Also, manual and automated DAWM and FWML volumes were not significantly different in both datasets. MTR values (expressed as mean and standard deviation, arbitrary units) were significantly lower in manually derived FWML compared with DAWM in both datasets: 1) FWML: 37.1 ± 3.2 vs DAWM: 43.3 ± 2.1; t=13.2; p<0.0001, and 2) FWML: 32.5 ± 3.9 vs DAWM: 38.8 ± 1.7; t=9.8; p<0.0001. Similar results were obtained using the automatic derived masks in both datasets: 1) FWML: 35.8 ± 2.8 vs DAWM: 43.1 ± 1.6; t=15.3; p<0.0001, and 2) FWML: 31.3 ± 3.1 vs DAWM: 38.1 ± 1.5; t=12.8; p<0.0001. MTR was also significantly lower in manually derived DAWM masks compared with normal appearing white matter (NAWM) in both datasets: 1) NAWM: 46.7 ± 1.3; t=10.1; p<0.0001, and 2) NAWM: 39.3 ± 0.8; t=3.1; p=0.003. Similar results were obtained using the automatic derived masks in both datasets: 1) NAWM: 46.3 ± 1.1; t=13.7; p<0.0001, and 2) NAWM: 39.9 ± 1.1; t=9.6; p<0.0001.

In both datasets, manually derived FWML and DAWM MTR values showed significant correlations with normalized FLAIR (r=−0.35 to −0.67; p=0.06 to <0.0001), T2w (r=−0.60 to −0.72; p=0.003 to <0.0001), T1w/T2w (r=0.63 to 0.77; p=0.002 to <0.0001), and T1w (r=0.77 to 0.92; p<0.0001) intensities. Finally, normalized intensity values obtained using automatic derived masks were significantly correlated with the manually derived values in both datasets.

**Conclusions:** The separation of FWML and DAWM on MRI scans of MS patients using automated intensity thresholds on T2w images is feasible. MTR values are significantly lower in FWML than DAWM, and DAWM values are significantly lower than NAWM, reflecting potentially greater demyelination within focal lesions. Normalized intensity values of different MRI contrasts exhibit a significant correlation with MTR values in both tissues of interest and could be used as a proxy to assess demyelination when MTR images are not available.

## 1. INTRODUCTION

Multiple sclerosis (MS) is an inflammatory, demyelinating, and neurodegenerative disease of the central nervous system (CNS) that affects the white matter (WM) and the grey matter (GM) (Compston and Coles 2008; Filippi et al. 2018). The characteristic lesions of MS are WM demyelinating plaques, with a variable number of inflammatory cells according to their degree of activity: acute lesions, chronic active lesions, chronic inactive lesions, remyelinating lesions or “shadow plaques” (Kutzelnigg and Lassmann 2014; Popescu, Pirko, and Lucchinetti 2013; Kuhlmann et al. 2017). These focal WM lesions (FWML) are relatively straightforward to detect and quantify on magnetic resonance imaging (MRI), where they appear as areas of high signal intensity on T2 weighted (T2w) and fluid attenuated recovery (FLAIR) images, and low signal intensity on T1 weighted (T1w) MRI contrasts (Reich, Lucchinetti, and Calabresi 2018). Additionally, a different type of T2w/FLAIR WM hyperintensity is also visible in the brain MRI of MS patients, which consists of mildly hyperintense areas of an intermediate value between that of normal-appearing WM (NAWM) and FWML (Seewann et al. 2009; Laule et al. 2013; Laule et al. 2011). This type of hyperintensity is referred to as diffusely abnormal white matter (DAWM) and presents as diffuse areas of increased T2w signal with poorly defined edges, in contrast to the sharp boundaries that characterized FWML (Laule et al. 2011) (Figure 01).

**Figure 1:**
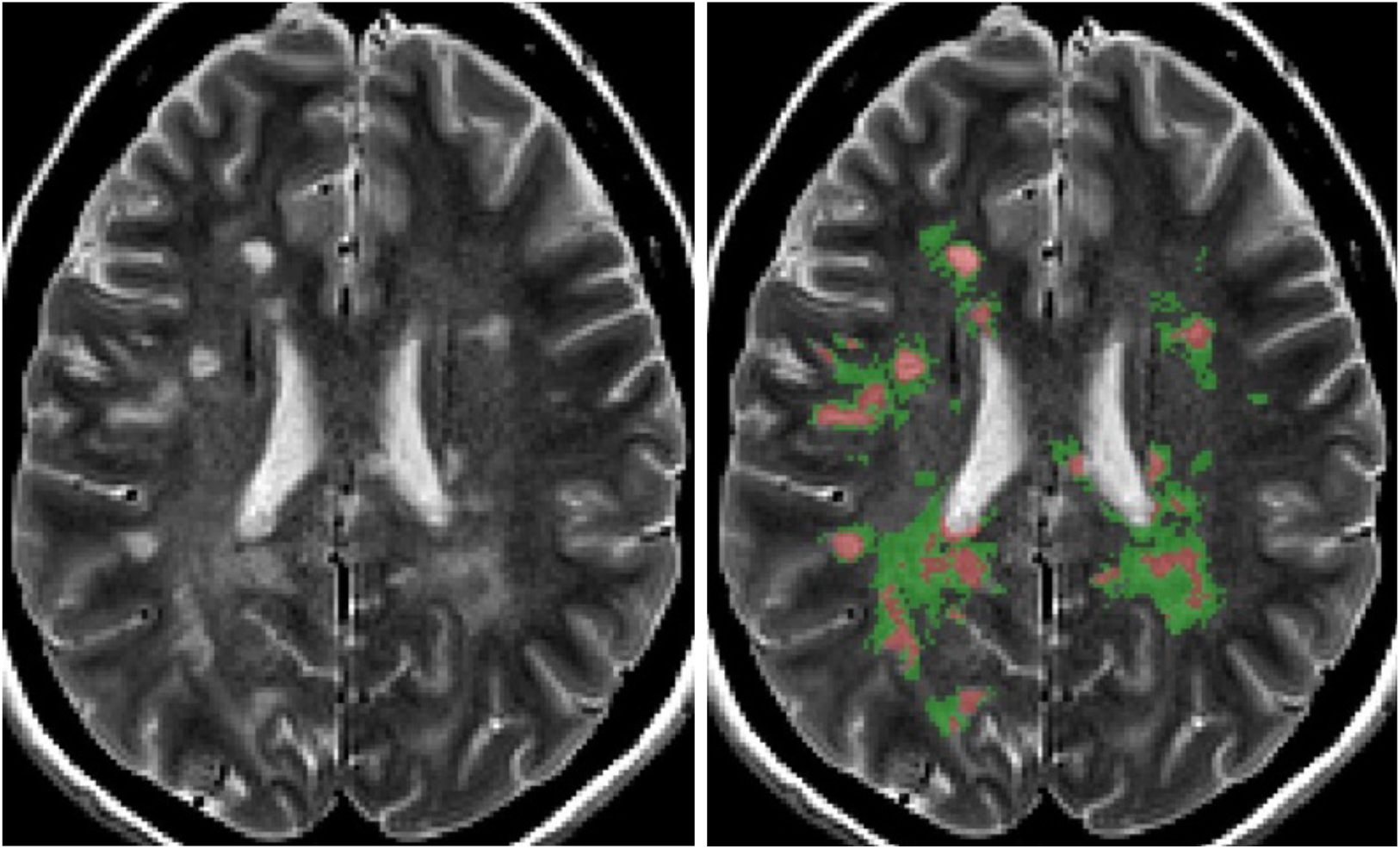
Example of manually derived masks for focal white matter lesions in red, and diffusely abnormal white matter hyperintensity in green.

Previous histopathology and MRI studies have addressed the differences between FWML and DAWM, showing quantitative changes in the degree of demyelination, axonal loss and immune cell density, which are all more severe in FWML (Seewann et al. 2009). MRI studies using semi-quantitative techniques, such as magnetization transfer ratio (MTR), which is sensitive to myelin content, have also shown significant differences in the values of FWML and DAWM (Ge et al. 2003). The relationship between the load of focal and diffuse lesions in the WM with specific MS clinical types (relapsing-remitting and progressive MS) has also been studied in the past, with results suggesting that diffuse changes may relate to progression and cortical pathology, while focal plaques would be characteristic of a relapsing-remitting course (Kutzelnigg et al. 2005). Interestingly, all of these previous studies have measured the areas of DAWM through fully manual segmentation done by imaging researchers. To our knowledge, there is no fully automated method for differentiating DAWM and focal lesions. Given that manually traced lesion masks are time-consuming to obtain and prone to inter-rater and intra-rater variability (Garcia-Lorenzo et al. 2013) an automated technique that consistently applies the same rule for separation of these tissues is highly advantageous.

Our study presents an MRI method that automatically separates DAWM from FWML and NAWM and that is comparable to the manual results of an imaging expert. Additionally, since this technique allows us to identify NAWM that is truly normal-appearing on conventional T2w and FLAIR imaging, we assess whether this “purer” NAWM could be used as a reference tissue for normalization of image intensity values on non-quantitative MRI contrasts: FLAIR, T2w, T1w/T2w ratio, and T1w. We then assess the correlation of these normalized values to MTR in the two hyperintense areas of interest: FWML and DAWM.

## 2. METHODS

### 2.1. Study Design

Forty MRI scans of two different MS cohorts were used in this study. The first dataset of 20 MRI scans was used to fulfill two goals: 1) to separate and qualitatively assess the separation of areas of FWML and DAWM using minimum and maximum manually-selected intensity-thresholds of DAWM, and 2) to automate the technique by regressing the manual intensity thresholds values on NAWM intensities obtained with automated tissue segmented masks. The second dataset of 20 MRI scans was used to replicate the findings.

### 2.2. Participants

#### Dataset 1

Twenty MS patients, 10 with relapsing-remitting MS (RRMS) and 10 with secondary progressive MS (SPMS), between 38 and 56 years of age, were recruited from the London, Ontario MS Clinic. The median Expanded Disability Status Scale (EDSS) score was 3 (range 1-6.5).

#### Dataset 2

Twenty MS patients, 13 RRMS and 7 SPMS between 19 and 69 years of age, were recruited from the Montreal Neurological Institute MS Clinic. The median EDSS score was 3 (range 0-7).

The demographic details of the participants and the differences across MS types (RRMS vs SPMS) are presented in Table 01.

**Table 01:** Demographic and clinical characteristics of the participants.

|  | **MS Patients** | **RRMS Patients** | **SPMS Patients** | **Significance** |
| --- | --- | --- | --- | --- |
| **Dataset 1 (London, Ontario)** |  |  |  |  |
| **Number** | 20 | 10 | 10 |  |
| **Male/Female** | 6/14 | 3/7 | 3/7 | 1 <sup>(1)</sup> |
| **EDSS** | 3 [1,6.5] | 1.5 [1,3] | 4.5 [3,6.5] | 0.001 <sup>(2)</sup> |
| **Age at Onset** | 32 ± 7.4 | 34 ± 8.22 | 30 ± 6.55 | 0.15 <sup>(3)</sup> |
| **Disease Duration** | 15 ± 7.87 | 12 ± 7.79 | 18 ± 7.22 | 0.05 <sup>(3)</sup> |
| **Dataset 2 (Montreal, Quebec)** |  |  |  |  |
| **Number** | 20 | 13 | 7 |  |
| **Male/Female** | 5/15 | 3/10 | 2/5 | 0.8 <sup>(1)</sup> |
| **EDSS** | 3 [0,7] | 2 [0,3.5] | 6 [3.5,7] | 0.0001 <sup>(2)</sup> |
| **Age at Onset** | 37 ± 9.7 | 34 ± 9.68 | 42 ± 8.10 | 0.04 <sup>(3)</sup> |
| **Disease Duration** | 15 ± 8.84 | 13 ± 8.54 | 20 ± 9.36 | 0.15 <sup>(3)</sup> |

Values are reported as Number, Mean ± SD or Median, [range]. Abbreviations: MS=Multiple sclerosis; RRMS=Relapsing-remitting multiple sclerosis; SPMS=Secondary progressive multiple sclerosis; SD=Standard deviation; EDSS=Expanded Disability Status Scale. (1) Chi-square test. (2) Mann-Whitney U test. (3) T-test.

The studies were approved by the Institutional Research Ethics Boards of the University of Western Ontario and McGill University. Each participant gave written informed consent.

### 2.3. MRI acquisition

Dataset 1 was acquired at the Centre for Functional and Metabolic Mapping at the Robarts Research Institute, University of Western Ontario, using a 3T Siemens Magnetom Prisma MRI scanner. Dataset 2 was acquired at the McConnell Brain Imaging Centre of the Montreal Neurological Institute, McGill University, also using a 3T Siemens Magnetom Prisma. The contrasts acquired in both sites were similar and included: 1) 3D-magnetization-prepared-2-rapid-acquisition-gradient-echoes (MP2RAGE) sequence, yielding a 3D T1w image and a quantitative T1-map 2) 3D FLAIR, 3) 2D dual-echo, turbo-spin-echo (TSE) yielding proton-density (PDw) and T2w images, and 4) 3D fast-low-angle-shot (FLASH) with and without an MT pulse, to compute MT ratio (MTR) images. The detailed acquisition parameters are listed in Table 02.

**Table 02:** MRI acquisition parameters.

|  | 3T T1w | 3T PDw/T2w | 3T FLAIR | 3T MT-on/off |
| --- | --- | --- | --- | --- |
| **Dataset 1 (London, Ontario)** |  |  |  |  |
| **Sequence** | 3D MP2RAGE | 2D-dual echo TSE | 3D TSE | 3D GRE |
| **Orientation** | Sagittal | Axial | Sagittal | Axial |
| **TR (ms)** | 5000 | 2350 | 6000 | 36 |
| **TE (ms)** | 2.98 | 22/87 | 356 | 3.86 |
| **TI (ms)** | 700 | n-a | 2200 | n-a |
| **Flip Angle** | 4 | 90 | 90 | 10 |
| **Slices** | 176 | 120 | 176 | 192 |
| **Voxel size (mm<sup>3</sup>)** | 1x1x1 | 1x1x1.5 | 1x1x1 | 1x1x1 |
| **Scan time (min:sec)** | 8:22 | 5:26 | 8:44 | 8:10 |
| **Dataset 2 (Montreal, Quebec)** |  |  |  |  |
| **Sequence** | 3D MP2RAGE | 2D-dual echo TSE | 3D TSE | 3D GRE |
| **Orientation** | Sagittal | Axial | Sagittal | Axial |
| **TR (ms)** | 5000 | 3000 | 6000 | 36 |
| **TE (ms)** | 2.76 | 11/99 | 356 | 4.92 |
| **TI (ms)** | 940 | n-a | 2200 | n-a |
| **Flip Angle** | 4 | 165 | 180 | 5 |
| **Slices** | 208 | 72 | 176 | 192 |
| **Voxel size (mm<sup>3</sup>)** | 1x1x1 | 1x1x2 | 1x1x1 | 1x1x1 |
| **Scan time (min:sec)** | 7:07 | 3:47 | 8:44 | 7:10 |

**Abbreviations:** TR= Repetition time; TE= Echo time; TI= Inversion time; T1w= T1-weighted; T2w= T2-weighted; PDw= Proton density weighted; FLAIR= Fluid attenuated inversion recovery; MT= Magnetization transfer; n-a= not applicable.

### 2.4. MRI analysis

#### 2.4.1. Image processing

All 3T MRI contrasts were co-registered in 3D, prior to lesion segmentation, using the following image processing pipeline: 1) brain mask extraction (Eskildsen et al. 2012), 2) bias field correction (Sled, Zijdenbos, and Evans 1998), 3) linear 9-parameter registration of the T1w image to standard MNI stereotactic space, International Consortium for Brain Mapping (ICBM) 152 model (Mazziotta et al. 2001) (using gradient orientations of minimal uncertainty) (Dadar, Fonov, and Collins 2018; De Nigris, Collins, and Arbel 2012), 4) 6 parameter rigid body registration of each of the other contrasts to the T1w image and 5) resampling of all contrasts to MNI stereotactic space via a single concatenated transform.

#### 2.4.2. Focal White Matter Lesions and Diffusely Abnormal White Matter Segmentation Using Manually Selected Thresholds

The segmentation of the two types of hyperintensities affecting the WM: FWML and DAWM, was done through the following steps:

1. An automated Bayesian classifier (Elliott 2016) was used to generate an automated WM lesion mask for each subject, which was corrected by an experienced rater (JM). This mask is referred as the Global WM Lesion (GWML) mask. GWML masks include FWML and areas of adjacent DAWM closer in intensity to FWML.
2. An experienced rater (JM) identified a minimum and a maximum intensity threshold for DAWM. The goal was to capture the highest DAWM intensity value that was lower than FWML intensity, and the lowest intensity value that was higher than NAWM intensity.
3. An experienced rater (JM) manually traced NAWM areas that appeared completely normal on T2w and FLAIR images in four brain regions: left and right frontal lobes and left and right temporal lobes. Each NAWM area was composed of 100 voxels. Mean intensity of NAWM was calculated using all 400 voxels.
4. The GWML mask was dilated by 3 voxels in 3D to capture additional areas of DAWM, potentially missed by the Bayesian classifier, due to their less hyperintense signal (areas closer to the minimum intensity threshold of DAWM). In this step, tissue masks for GM, WM, and cerebrospinal fluid (CSF), based on a Multi-Atlas Label Fusion method (MALF) (Sabuncu et al. 2010) were used to limit the dilation of the lesion mask, to avoid the inclusion of cortical GM or CSF as part of the dilated GWML mask.
5. The dilated GWML mask was separated into two classes: FWML, using the maximum DAWM intensity threshold to exclude the DAWM areas, and DAWM, using the minimum DAWM intensity threshold or the NAWM mean intensity value, if the latter was higher than the former, to exclude NAWM.
6. The final FWML and DAWM masks were reviewed by an experienced rater (JM) to qualitatively assess and assure the correct separation of the two intensity classes.

Figure 1 shows an example of the FWML and DAWM masks for a subject using manually-selected intensity thresholds

#### 2.4.3. Automation of Focal White Matter Lesions and Diffusely Abnormal White Matter Segmentation

1. To calculate the minimum and maximum thresholds automatically, the following two linear regressions models were fit: Tmin = Wmin * NAWMm Tmax = Wmax* NAWMm Where Wmin and Wmax were the weights of the model estimated by the linear regression, Tmin and Tmax were the manually selected thresholds, and NAWMm was the mode of the intensity histogram of the NAWM tissue in the T2w image obtained from the MALF tissue segmentation (Sabuncu et al 2010). After estimating Wmin and Wmax, we were able to calculate automatic Tmin and Tmax thresholds and separate FWML and DAWM voxels in the data.
2. The automated thresholds were applied to dataset 1 and 2 to generate automated FWML and DAWM masks for each case.
3. The automated FWML and DAWM masks were reviewed by an experienced rater (J.M.) to qualitatively assess and assure the correct separation of the two intensity classes, and to qualitatively compare to the manually obtained masks.

Figure 2 shows and example of manually derived masks (second row) compared to the automatic-derived masks (third row) of FWML and DAWM.

**Figure 2:**
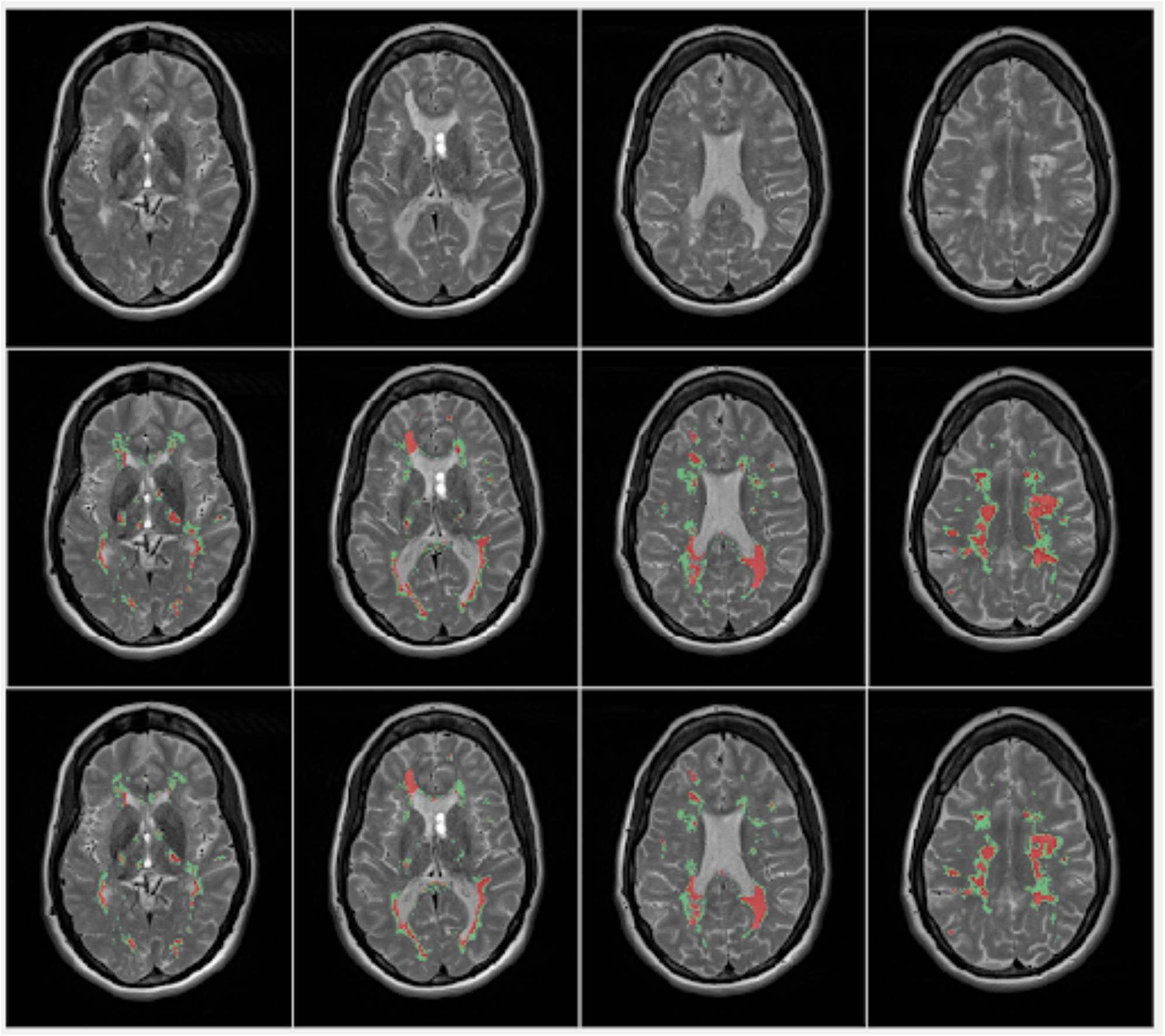
Example of manually derived masks (second row) and automatically derived masks (third row) for focal white matter lesions in red, and diffusely abnormal white matter hyperintensity in green.

#### 2.4.4. Calculation of Normalized Intensity Values

MRI intensity values are not an absolute scale, so they need to be normalized for processing. We used two approaches to generate normalized intensity values:

1. The mean intensity value within the manual NAWM masks was calculated for all MRI contrasts of interest: T2w, T1w, and FLAIR. We called this value the NAWM-Normalization Factor (NAWM-NF). The ratio of the mean intensity of FWML (or DAWM) to the NAWM-NF was calculated for each corresponding MRI contrast.
2. Automated NAWM normalization factor (A-NAWM-NF): We used automatic MALF WM masks to which we subtracted: 1) the DAWM and FWML masks, 2) MALF cortical GM and deep GM masks dilated by one voxel, and 3) MALF ventricular masks dilated by 1 voxel. The subtractions from the WM masks were done to obtain a NAWM mask that excluded not only lesional voxels, but also partial volume voxels with other tissue types (i.e. CSF). The mean on the intensities under this automatic NAWM mask was used as the A-NAWM-NF.

#### 2.4.5 Calculation of MTR values

MT-on and MT-off contrasts were used to produce MT ratio images (MTR). Mean MTR values were obtained for FWML and DAWM masks, both manually and automatically-derived. The goal was to assess the relationship between the MTR values and normalized-intensity values, because post-mortem analysis of MS lesions has shown that MTR is sensitive to the degree of demyelination (Schmierer et al. 2004; Chen et al. 2007).

#### 2.4.6. Calculation of T1w/T2w ratio

The ratio of normalized T1w and T2w values was calculated within the DAWM and FWML masks, again manually and automatically-derived values. This approach has been used to reduce the image intensity bias and enhance the signal to noise ratio for myelin. (Glasser and Van Essen 2011)

### 2.5. Statistical analysis

Subject demographic data was compared between datasets using a chi-square test, a T-test or a Mann-Whitney-U test. MRI measurements (i.e. volumes and MTR values) were compared between MS types using a T-Test or a Mann-Whitney-U test, as appropriate given the normal or non-normal distribution of the variables. FWML and DAWM volumes manually and automatically generated were compared using Dice-kappa, to assess spatial overlap of the masks, and using Paired-T-test or Wilcoxon Signed-rank test, according to their distribution, to assess differences of the volumetric values. MTR values in areas of NAWM, FWML and DAWM were compared with a T-test. The relationship between MTR values and normalized intensity values was evaluated with Pearson correlation. All statistical analyses were performed using MATLAB R2018a and SPSS v.24.

### 2.6 Data and code availability statement

The extracted numerical MRI data and code that support the results of this study are available from the corresponding author upon request.

## 3. RESULTS

### 3.1. Study Population Features

The proportions of MS course (RRMS or SPMS) and sex across datasets were not significantly different. The age of the participants at onset and the duration of the disease at the time of the MRI scan were not significantly different across datasets (Table 01).

### 3.2. Automatic Minimum and Maximum Intensity Thresholds of Diffusely Abnormal White Matter

Based on dataset 1, the estimated values for Wmin and Wmax were 1.15 and 1.39, respectively. Therefore, FWML masks were generated with all voxels within the GWML mask higher than 1.39 × MALF NAWM mode value.

DAWM masks were generated using any voxel within the GWML mask between 1.15 and 1.39 × MALF NAWM mode value.

### 3.3. Diffusely Abnormal White Matter Intensity Thresholds and Normal Appearing White Matter Normalization Factor

We assessed the correlation between the DAWM minimum and maximum thresholds and the NAWM-NF and A-NAWM-NF to evaluate the consistency of the individual values selected by the rater. The correlation between the minimum and maximum DAWM thresholds were r=0.95 (p<0.0001) in dataset 1 and r=0.90 (p<0.0001) in dataset 2. The correlations between the minimum DAWM threshold and the NAWM-NF were r=0.92 (p<0.0001) in dataset 1 and r=0.91 (p<0.0001) in dataset 2. Finally, the correlations between the maximum DAWM threshold and the NAWM-NF were r=0.88 (p<0.0001) in dataset 1 and r=0.90 (p<0.0001) in dataset 2. The correlations between the minimum DAWM threshold and the A-NAWM-NF were r=0.95 (p<0.0001) in dataset 1 and r=0.93 (p<0.0001) in dataset 2. Finally, the correlations between the maximum DAWM threshold and the NAWM-NF were r=0.96 (p<0.0001) in dataset 1 and r=0.92 (p<0.0001) in dataset 2.

The consistent and linear relationship of the thresholds and the NAWM values was the rationale to use a regression approach for the automation of the thresholds.

### 3.4. Focal White Matter Lesion and Diffusely Abnormal White Matter Volumes

**3.4.1.** Dice-kappa values were calculated to assess spatial overlap of FWML and DAWM masks obtained manually and automatically, with results ranging from 0 to 1: 0 indicating lack of spatial overlap and 1 perfect overlap of all voxels. The mean Dice-kappa values for dataset 1 were: 0.8 for DAWM masks and 0.7 for FWML masks. In dataset 2, mean dice-kappa values were: 0.8 for DAWM and 0.8 for FWML.

**3.4.2** Manually derived FWML and DAWM volumes in dataset 1 were not significantly different than automated FWML and DAWM volumes: manual FWML volume 9.0cc ± 9.8 and automated FWML volume 9.5cc ± 9.6 (p=0.7); manual DAWM 15.8cc ± 12.1 and automated DAWM 15.3cc ± 8.5 (p=0.8). Similar results were obtained in dataset 2: manual FWML 4.3cc ± 4.8 and automated FWML 5.1cc ± 7.4 (p=0.5); manual DAWM 24.2cc ± 16.6 and automated DAWM 22.5cc ± 22.0 (p=0.5).

The manual volumes of FWML and DAWM were assessed across MS types: RRMS and SPMS, in both datasets. We found significantly higher DAWM volume in SPMS participants of dataset 1 (p=0.02), but not in dataset 2. The volumes of FWML of dataset 1 and 2 were not significantly different. See detailed results in Table 3.

**Table 03:** FWML and DAWM Volumes expressed in cubic centimetres.

|  | MS type | Mean | SD | Median | Range | Significance |
| --- | --- | --- | --- | --- | --- | --- |
| Dataset 1 (London, Ontario) |  |  |  |  |  |  |
| FWML Volume | RRMS | 5.95 | 5.23 | 4.54 | 0.62-8.09 | 0.16 <sup>(1)</sup> |
|  | SPMS | 11.70 | 11.65 | 6.48 | 1.73-33.37 |  |
| DAWM Volume | RRMS | 9.71 | 6.71 | 7.90 | 1.34-22.34 | **0.02** <sup>(2)</sup> |
|  | SPMS | 20.16 | 13.65 | 19.88 | 4.03-43.10 |  |
| Dataset 2 (Montreal, Quebec) |  |  |  |  |  |  |
| FWML Volume | RRMS | 3.53 | 3.48 | 2.53 | 0.33-10.83 | 0.16 <sup>(1)</sup> |
|  | SPMS | 6.05 | 7.05 | 4.46 | 0.60-20.74 |  |
| DAWM Volume | RRMS | 19.20 | 16.68 | 19.15 | 2.60-62.90 | 0.22 <sup>(1)</sup> |
|  | SPMS | 21.94 | 23.52 | 14.87 | 1.47-68.26 |  |

(1) Mann-Whitney-U test; (2) T-test

### 3.5. Focal White Matter Lesion and Diffusely Abnormal White Matter MTR values

MTR values were significantly lower in manually derived FWML compared with DAWM in both datasets. In dataset 1 the MTR mean and SD values in FWML were 37.1 ± 3.2 vs 43.3 ± 2.1 in DAWM (p<0.0001). In dataset 2 FWML MTR values were 32.5 ± 3.9 vs 38.0 ± 1.7 in DAWM (p<0.0001). Additionally, MTR was significantly lower in DAWM compared with NAWM in both datasets: 1) NAWM: 46.7 ± 1.3 (p<0.0001), and 2) NAWM: 39.3 ±0.8 (p=0.003).

Similar results were obtained when assessing MTR values using automated derived FWML and DAWM masks in both datasets: Dataset 1: FWML: 35.8 ± 2.8 vs DAWM: 43.1 ± 1.6; t=15.3; p<0.0001. Dataset 2: FWML: 31.3 ± 3.1 vs DAWM: 38.1 ± 1.5; t=12.8; p<0.0001. Finally, MTR values obtained using automated DAWM were also significantly lower than MALF generated NAWM in both datasets: dataset 1: 46.3 ± 1.1; t=13.7; p<0.0001, and in 2: NAWM: 39.9 ± 1.1; t=9.6; p<0.0001. (Figure 3).

**Figure 3:**
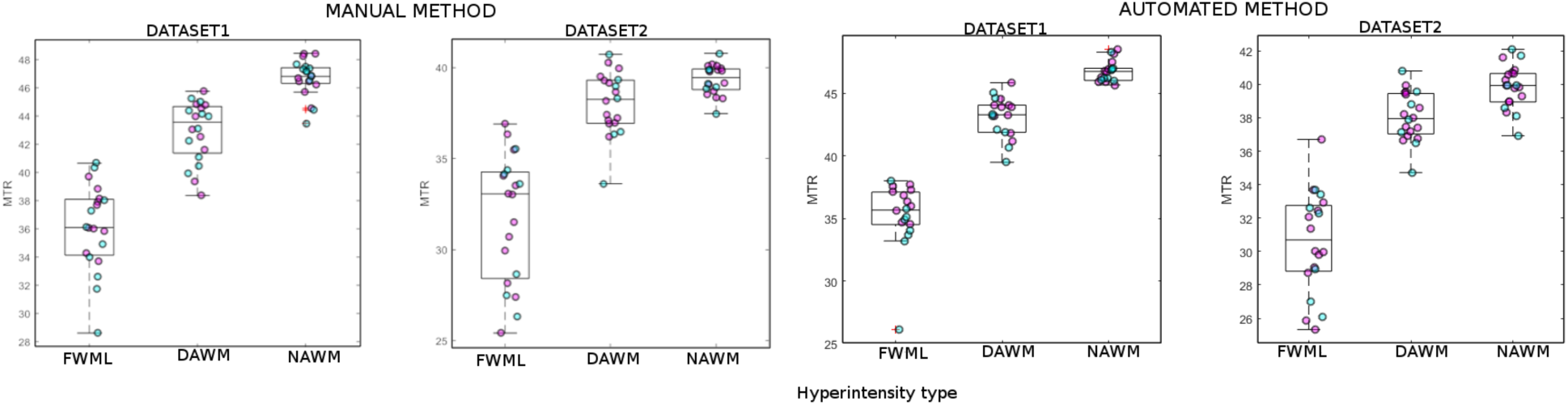
Mean MTR values in manually (first two panels) and automatically (last two panels) derived FWML, DAWM, and NAWM masks in Dataset 1 and Dataset 2. FWML: Focal White Matter Lesion; DAWM: Diffusely Abnormal White Matter. Magenta: relapsing-remitting MS; Cyan: secondary progressive MS.

### 3.6. MTR and normalized intensity values

MTR values showed significant correlations with normalized FLAIR (p<0.06), T2w (p<0.003), T1w/T2w (p<0.001), and T1w (p<0.0001) intensities in manually derived FWML and DAWM in both datasets, using NAWM-NF (Figures 4 and 5). Note that since the FWML and DAWM regions are hyperintense compared with NAWM on T2w and FLAIR images and hypointense in T1w images, the normalized T2w and FLAIR values are greater than 1.0 and the normalized T1w values are smaller than 1.0. Since normalized T1w values are lower than 1.0 and T2w values are higher than 1.0, the T1w/T2w ratio values are also lower than one.

**Figure 5:**
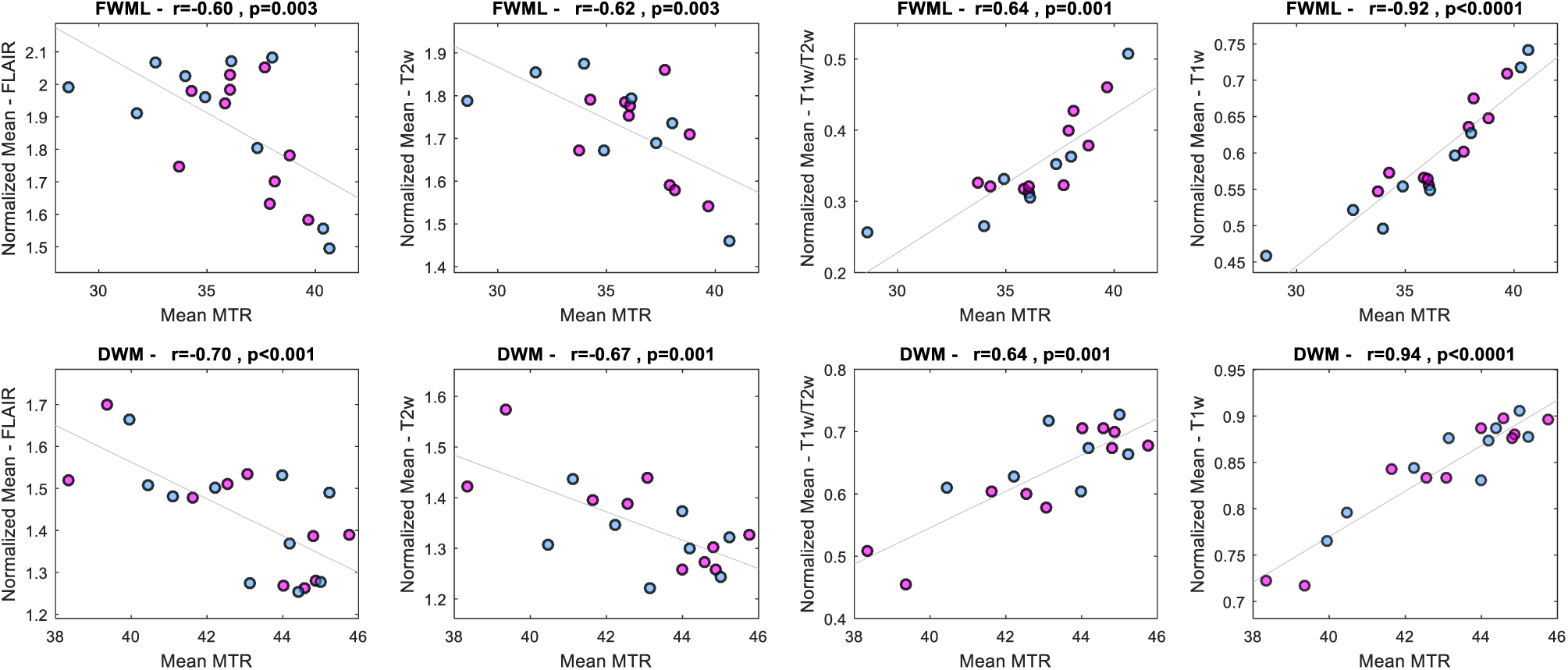
Association of Mean MTR values and FLAIR, T2w, T1w/T2w, and T1w intensity normalized values of dataset 1. FLAIR: fluid attenuated inversion recovery; T2w: T2 weighted, T1w: T1 weighted; Magenta: relapsing-remitting MS; Cyan: secondary progressive MS.

**Figure 06:**
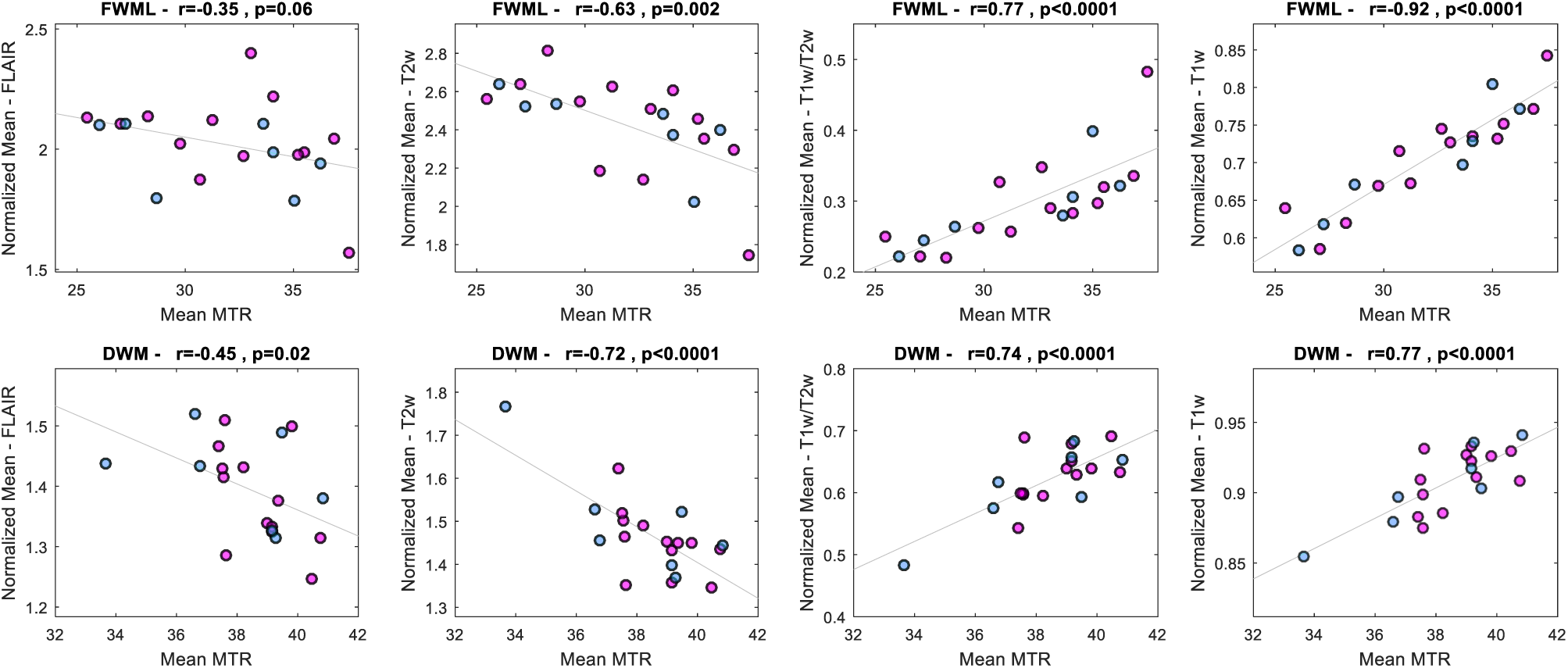
Association of Mean MTR values and FLAIR, T2w, T1w/T2w, and T1w intensity normalized values of dataset 2. FLAIR: fluid attenuated inversion recovery; T2w: T2 weighted, T1w: T1 weighted; 1: relapsing-remitting MS; 2: secondary progressive MS.

Finally, all normalized intensity values obtained using manually derived FWML and DAWM masks showed significant correlation with the normalized values obtained using automated masks and the A-NAWM-NF in both datasets: FLAIR: 0.5<r<0.7 (p<0.02), T2w 0.4<r<0.7 (p<0.6), T1w/T2w 0.5<r<0.8 (p<0.02), T1w 0.7<r<0.8 (p<0.002).

## 4. DISCUSSION

Our automated T2w intensity thresholding technique to separate FWML from DAWM was tested in two MRI datasets, acquired in different centers with similar 3T MRI acquisition protocols. Our results were consistent at various levels across datasets.

First, the qualitative assessment of each scan allowed the verification of FWML as sharply delimited areas of hyperintensity, while DAWM areas were diffuse regions of intermediate brightness between that of FWML and NAWM, with poorly defined edges (Figure 2).

Second, the quantitative assessment of MTR values in FWML and DAWM in both datasets consistently showed significantly lower values in FWML (p<0.0001), indicating more severe demyelination in focal lesions (Figure 4 and 5), as expected. This supports our technique as a suitable separation tool to differentiate two distinct types of tissue damage. Our finding is in line with previous histopathology-MRI studies, showing that the areas of DAWM represent regions of decreased myelin phospholipids in histology and reduced myelin water fraction in MRI (Laule et al. 2011; Laule et al. 2013). Another study correlated various histological quantifications (i.e. myelin content, axonal loss) in NAWM, DAWM, and FWML with different quantitative MRI measurements (i.e. T1 and T2 relaxation times and MTR values), finding significant differences in axonal and myelin content, progressively decreasing from NAWM, to DAWM, and to FWML (Seewann et al. 2009). These changes paralleled the MRI quantitative findings of the same areas in terms of progressive increase of T1 and T2 relaxation times and progressive decrease of MTR values (Seewann et al. 2009).

Our third level of analysis in both datasets estimated the correlations between MTR and intensity normalized values on FLAIR, T2w, T1w/T2w, and T1w in areas of DAWM and FWML. We found, once again, consistent and significant correlations, showing in both groups a more robust performance (higher correlations with MTR values) of T1w normalized values, followed by T1w/T2w, T2w, and FLAIR normalized values. FLAIR normalized intensity values had poorer correlations with MTR. We attribute this behaviour to the change in the intensity of chronic lesions that evolve to a more severe destruction of the brain parenchyma, also known as chronic black hole evolution. At this stage, MS lesions start changing the characteristic hypersignal on FLAIR, and evolve slowly towards a lower signal typical of black holes. In other words, FWML on FLAIR are hyperintense as long as the level of tissue destruction has not increased the T1 to levels approaching those of CSF, when the signal would begin to be nulled by the inversion pulse. This signal change does not occur on T2w, where all MS lesions always show as hyperintense, even at the most chronic phase of black hole evolution, explaining the better correlation with MTR values than normalized FLAIR (Figure 8). However, the relationship of both normalized T2w and normalized FLAIR with MTR is modest, while that of normalized T1w and T1w/T2w are stronger, suggesting that normalized T1w and T1w/T2w could be used as myelin-sensitive contrasts in datasets where MTR or other more specific contrasts are not available.

**Figure 8:**
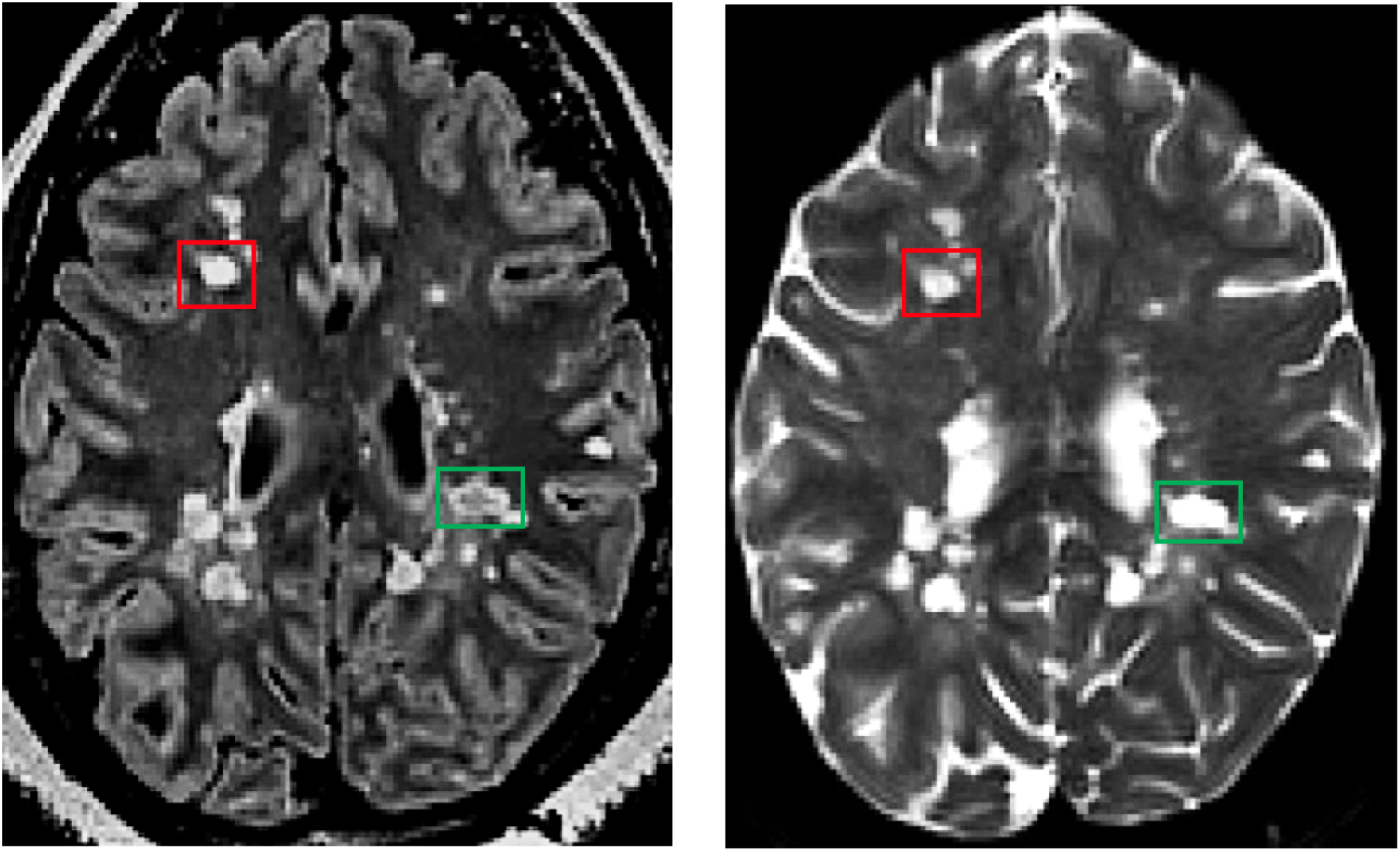
Comparison of chronic FWMLs in FLAIR and T2w images in one subject. FWML: Focal white matter lesions; FLAIR: Fluid attenuated inversion recovery; T2w: T2-weighted

To assess the consistency of the values manually selected by the expert (JM), we correlated the DAWM intensity thresholds and the NAWM-NF values (Figure 3). We found a linear relationship indicated by the high correlations (0.88<r<0.95; p<0.0001) across thresholds and NAWM-NF. This stable relationship of the values across scans and datasets was the rationale to automate the selection of threshold values by regressing them and determining the relationship with the NAWM histogram of intensities.

Our last assessment looked into the differences of DAWM volume and FWML volume across MS types (RRMS and SPMS) in both datasets. In dataset 1 we found a significantly higher volume of DAWM in SPMS compared to RRMS. This highlights the potential use of DAWM volume as a variable of interest to characterize SPMS cases. This finding did not reach significance in dataset 2, which had fewer SPMS cases than dataset 1 (7 as opposed to 10) and a non-normal distribution of DAWM volumes. This limitation in our current analysis could be addressed by using a larger sample size in future studies.

Another limitation to the proposed technique is that its starting point relies on the GWML mask. The rationale for using this mask as our initial starting point was to avoid the inclusion of vascular spaces, which are more frequently enlarged in MS (Achiron and Faibel 2002). These enlarged vascular spaces have intensity characteristics similar to FWML, hence using an overall WM mask as the starting point would have the disadvantage of including such regions as false positives. We recognize that if there are regions of DAWM that are farther than 3mm away from all voxels of the initial GWML mask, they would not be captured by our dilation, becoming false negatives not captured as DAWM. However, we did not encounter any examples of this type of false negative in our visual qualitative assessments, since the initial GWML mask already captures part of the DAWM as lesional tissue.

Finally, we would like to note the nomenclature that we have used to describe the different hyperintensities visualized on MRI. We used the term DAWM to describe ill-defined areas of hyperintensities visible on T2w and FLAIR. The diffuse nature of the abnormality only refers to these two MRI contrasts and should not be applied to other MRI techniques (i.e. MTR or DWI) (Laule et al. 2013), because numerous studies have shown that the NAWM in general is diffusely abnormal on quantitative MRI and histology (Moll et al. 2011).

## 5. CONCLUSIONS

The classification of FWML and DAWM on MRI scans of MS patients is feasible using automatically-selected intensity thresholds on T2w MRI. MTR values are significantly lower in FWML than DAWM, reflecting greater demyelination within focal lesions. Additionally, normalized intensity values of T1w and T1/T2 exhibit strong correlations with MTR values, both in DAWM areas and FWML, hence they could be utilized to assess demyelination when myelin-sensitive images such as MTR are not available.

